# igv-reports: Embedding interactive genomic visualizations in HTML reports to aid variant review

**DOI:** 10.1101/2025.10.29.685397

**Authors:** James T. Robinson, Helga Thorvaldsdottir, Jill P. Mesirov

## Abstract

We present igv-reports, a command-line tool to create standalone HTML pages embedding interactive genomic visualizations of read alignments and associated annotations to support variant inspection workflows. The reports contain all data and code required for visualization of the variant sites, with no dependencies on the input data files.

## Introduction

The accuracy of sequencing technologies and the tools that identify genomic variants based on the sequencing data has greatly improved over the years. However, incorrect variant calls may still occur due to sequencing anomalies and other causes, and expert human review of the underlying data can help identify false positives and validate correctly called variants. Therefore, visual inspection of read alignments in a genome browser is an important step in many sequencing workflows [1-3] such as validating variants in clinical settings using BAM or CRAM files. To support this a number of desktop and web applications have been developed or modified to support alignment viewing. Tools like the Integrative Genomics Viewer (IGV) [4,5] and web-based browsers such as JBrowse [6] and the UCSC Genome Browser [7] are powerful and widely used. However, they all require direct access to underlying data files during operation, which can complicate or prevent the sharing of the variant visualizations. To address this, tools have been developed to create static screenshots or precomputed images for reports or downstream analysis, e.g., IGV snapshot automator [8] and Alignoth [9]. However, this approach has a significant limitation: the visualization options are fixed at the time of image creation. Variant reviewers lack the ability to dynamically adjust coloring, sorting, grouping, and other visualization settings in response to specific characteristics, which would significantly enhance variant interpretation.

To address this, we present igv-reports, a tool that creates self-contained HTML reports for variant review by embedding the popular IGV plugin igv.js [10]. Unlike static screenshots, igv-reports supports the extensive visualization options developed for IGV, aiding in variant validation and interpretation. Once generated, these report files no longer depend on the original data files, making them easy to open as local files, share via email, post as static web pages, or integrate as output into variant pipelines.

## Features

The igv-reports tool takes as input (1) a list of genomic sites, specified as a file in tab-delimited format or one of several common genomic data file formats; (2) the reference genome, specified as a genome identifier or as a file that contains the reference sequence; and (3) one or more track files containing alignments, annotations, and other genomic data. Accepted file formats are listed in the *Implementation* section below. Input files can be loaded from the local file system or remotely by URL. Output is an HTML report file containing a table of variants and an igv.js genomic view of the tracks for each variant. The report file can be opened with a web browser from the local file system or shared on a server or web portal. Importantly, no access to the original input files is required to view and interact with the report. This is possible because all data for the regions of interest are embedded in the report file.

An example report is illustrated in Figure 1. The report was created with the following command, using test data from the igv-reports GitHub repository.

~~~
create_report test/data/variants/variants.vcf.gz --genome hg38
--info-columns GENE TISSUE TUMOR COSMIC_ID GENE SOMATIC
--tracks test/data/variants/variants.vcf.gz test/data/variants/recalibrated.bam
--output example_vcf.html
~~~

**Figure 1.**
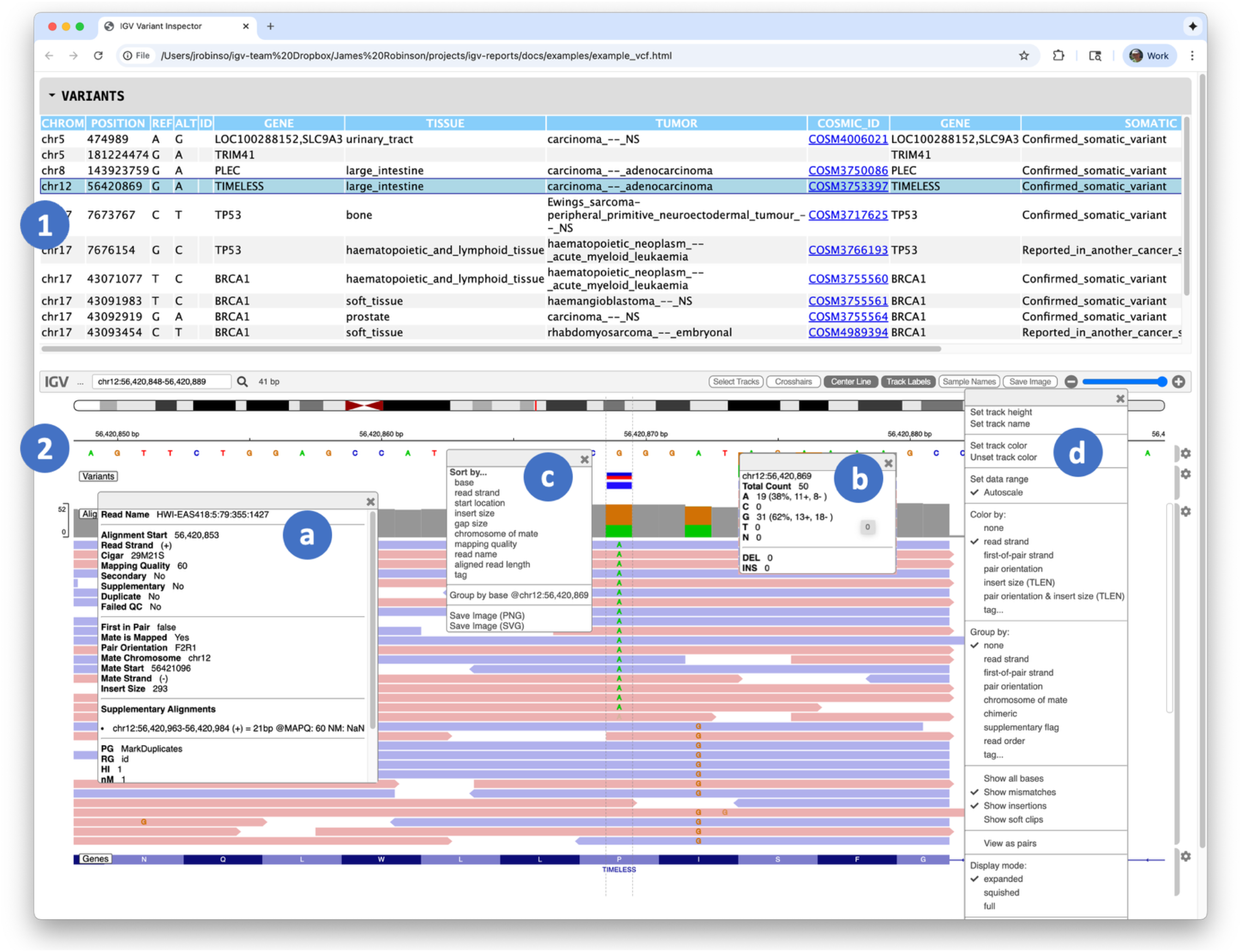
A variant report containing two main sections: **(1)** a table view of variants sites, and **(2)** an igv.js genomic visualization of variants and associated alignments. Overlaid on the igv.js viewer are four popups associated with the alignment track. Clicking on an element in the track displays further details, including metadata for a particular alignment **(a)**, and coverage details for the variant base **(b)**. The track has two menus: a context menu to control sorting of the alignments **(c)**, and a track configuration menu **(d)**. The interactive version of this report can be accessed at https://igvteam.github.io/igv-reports/examples/example_vcf.html.

The report includes a collapsible table view of variants coupled to a genomic view of supporting read alignments. Clicking on a variant in the table will display the tracks in the igv.js viewer, with the genomic view centered on the selected variant, including flanking regions to either side. The size of the flanking region is a user option, defaulting to 1 kilobase. The displayed tracks can be explored interactively using the igv.js track configuration and context menus, for example, to change coloring or sorting schemes.

Details of individual alignments and other displayed objects are available as popup text.

igv-reports offers multi-locus views for structural variants too large for a single display. This feature allows users to input two regions that encompass the variant endpoints. Multi-locus views are also employed for variants exceeding a length threshold specified as a command-line argument. See Supplementary Figure 1 for an example.

Several command-line options are available to customize a report. These include, for example, options to customize the variant table columns, include custom headers and footers, filter and sort alignments by various attributes, and adjust the size of the variant flanking region. Additionally, track options are fully customizable using igv.js track configuration objects. Finally, the report itself can be customized by supplying an alternative HTML template. Templates are standard HTML with placeholder tokens for the variant table data and dictionary of encoded igv.js sessions. In addition to the standard variant template, illustrated in Figure 1, igv-reports provides alternative templates in the GitHub repository. They include a template with filterable columns in the variant table (see Supplementary Figure 2), as well as templates for supporting output from the Trinity Cancer Transcriptome Analysis Toolkit (CTAT) gene fusion [11] and splice junction tools (see Supplementary Figures 3 and 4). Users can also provide their own custom templates.

The only tool we are aware of with a comparable set of features is jigv [12]. This tool, written in Nimble, takes a similar approach of creating a standalone HTML report containing interactive igv.js track views in the regions of a pre-defined list of variant sites, but augments it with special support for working with family pedigrees. While similar in design and operation, igv-reports offers some advantages when creating reports not involving pedigrees. First, jigv reports do not include a table view which means there is no annotated variant summary, and navigation is limited to clicking on next and previous buttons or the arrow keys to move between variants. Additionally, igv-reports offers more extensive customization options and supports a broader array of file formats. Finally, jigv does not support multi-locus views and therefore cannot handle the display of large structural variants.

## Implementation

The igv-reports tool is a command-line application written in Python. Supported input file formats include common genomic data formats: FASTA and TwoBit for the reference sequence; VCF, BED, MAF, BEDPE, and tab-delimited files for the list of variant sites; and BAM, CRAM, VCF, BED, GFF, MAF, and BEDPE for the track files. Processing proceeds as a loop through variant sites. For each site, metadata is extracted from the variant file for populating the corresponding row in the variant selection table. This data is stored in a TABLE_JSON dictionary. Next, data is extracted from the reference sequence file and track files for a genomic region centered on the variant site. For fast processing, data extraction uses the Python packages pysam [13] and py2bit [14] which take advantage of file indexing. Indexes are required for sequence and alignment formats, and they are recommended for all large files. The extracted data slices are then compressed and base64 encoded to form a data URI. The data URIs for each variant are incorporated in an igv.js session object stored in the SESSION_DICTIONARY object. Finally, the TABLE_JSON and SESSION_DICTIONARY objects are converted to JSON and injected into the HTML template at the corresponding token locations in the HTML template.

## Limitations

igv-reports is designed to generate self-contained HTML reports for the visual inspection and validation of variant sites. However, because all genomic data is embedded directly within the HTML file, the tool is not ideal for extremely large sets of variants. Our testing has shown that igv-reports is suitable for producing reports for variant sets ranging from hundreds up to 1,000 distinct variant sites. The exact limits on the number of variants can vary depending on factors such as the number of visualized tracks, the average depth of coverage in alignment tracks, the amount of metadata associated with variants and alignments, and the available memory on the client computer viewing the reports. A benchmark with an alignment file from the 1000 Genomes Project [15] of approximately 50x coverage produced a report of about 100 MB for 1,000 variants. While viewable on most modern web browsers, this is considered near the upper limit. For larger datasets, the report could be split into multiple files; although 1,000 variants may already surpass the limit of practical usefulness for visual validation.

## Availability

igv-reports is freely available under an MIT open source license. The Python package can be installed from PyPI at https://pypi.org. Source code, installation instructions, user documentation, and examples are available at https://github.com/igvteam/igv-reports.

## Acknowledgements

This work was supported by the National Institutes of Health [U24CA258406] to JPM and JR.

