## Supplemental Figures for "igv-reports: Embedding interactive genomic visualizations in HTML reports to aid variant review"

Note: All command lines run from the root directory of the igv repository: https://github.com/igvteam/igv-reports

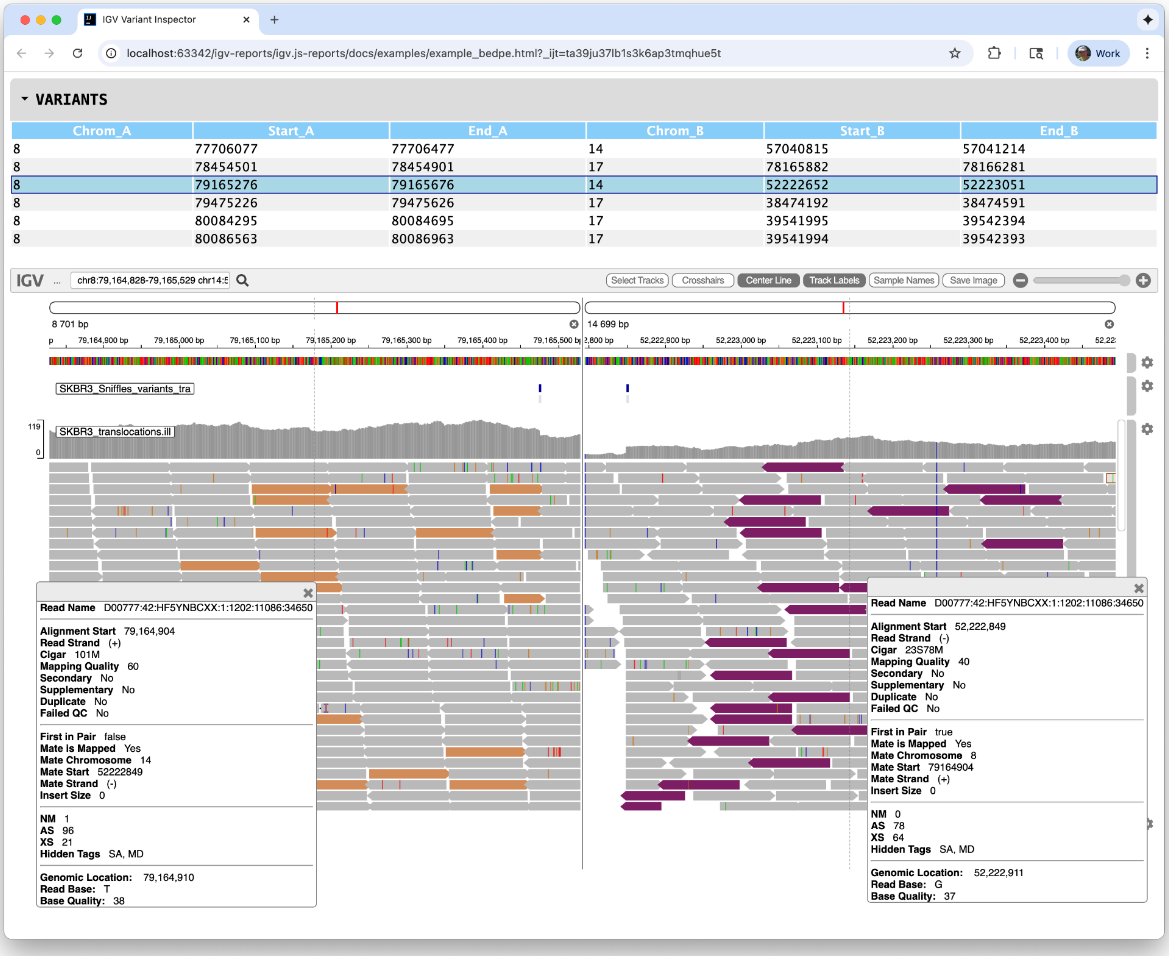


**Supplementary Figure 1.** Multi-locus view report. The selected variant represents an inter-chromosomal rearrangement between chromosomes 8 and 14. Alignments are colored by the chromosome of their mate. For an interactive version of this report see https://igvteam.github.io/igv-reports/examples/example_bedpe.html.

### Command line

```
create_reports test/data/variants/SKBR3_Sniffles_tra.bedpe
  -genome hg19
  -flanking 1000
  -tracks test/data/variants/SKBR3_Sniffles_variants_tra.vcf
  test/data/variants/SKBR3_translocations.ill.bam
  -output example_bedpe.html
```

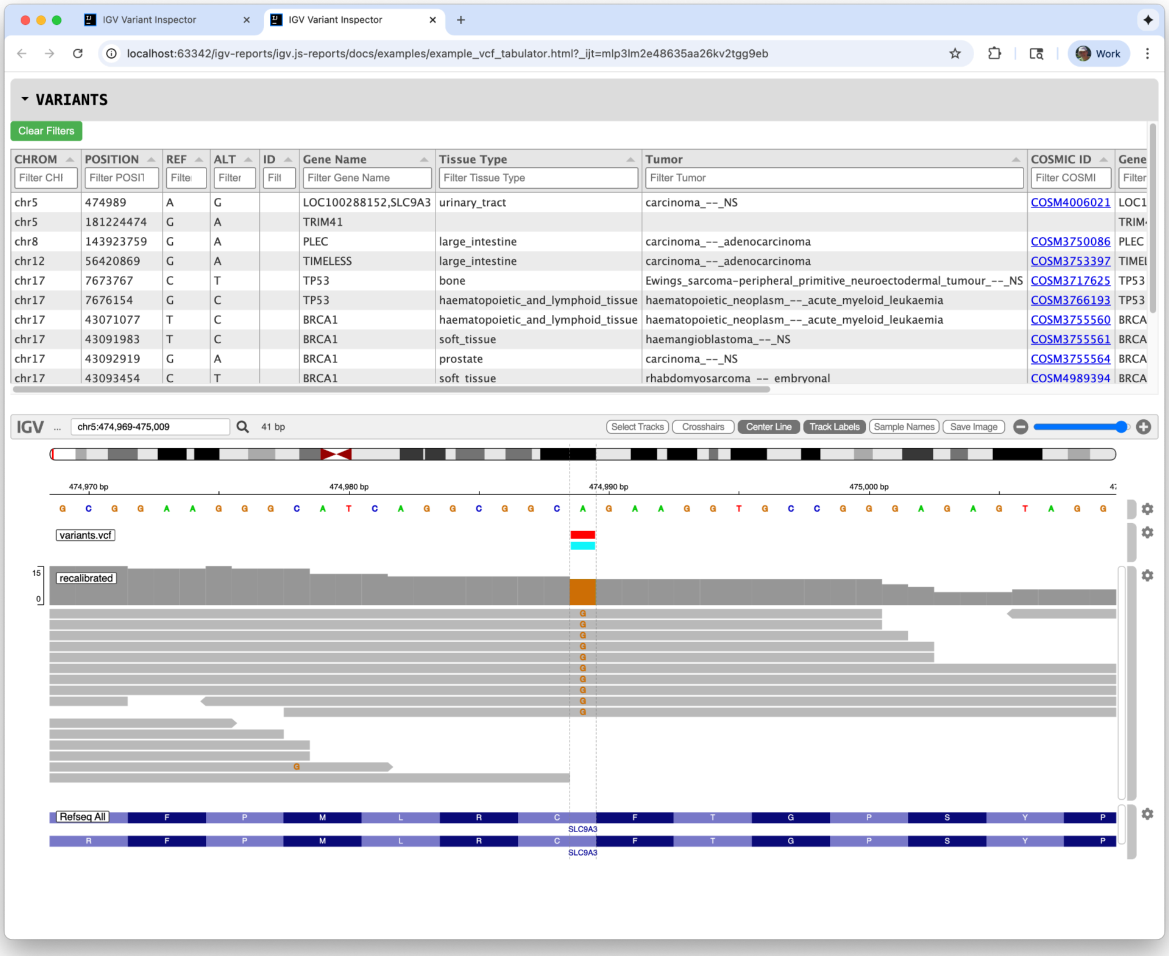


**Supplementary Figure 2.** Example of a report created with the "tabulator" template. This template allows filtering rows based on values for one or more columns. For an interactive version of this report see https://igvteam.github.io/igv-reports/examples/example_vcf_tabulator.html

### Command line

```
create_reports test/data/variants/variants.vcf.gz
  -genome hg38
  -ideogram test/data/hg38/cytoBandIdeo.txt
  -flanking 1000
  -info-columns GENE TISSUE TUMOR COSMIC_ID GENE SOMATIC
  -samples reads_1_fastq
  -sample-columns DP GQ
  -tracks test/data/variants/variants.vcf.gz test/data/variants/recalibrated.bam
  -tabulator
  -filter-config test/data/variants/filter_config.yaml
  -output example_vcf_tabulator.html
```

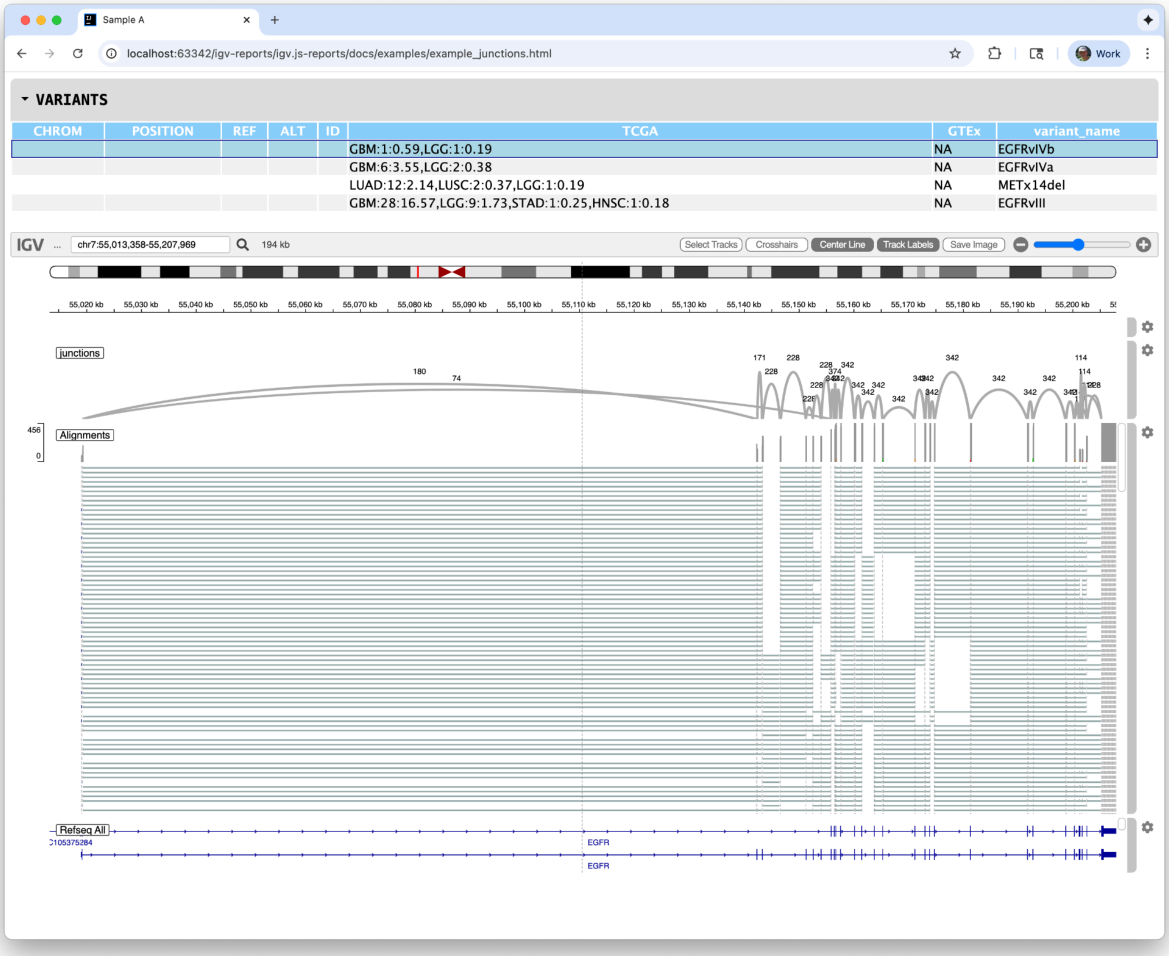


**Supplementary Figure 3.** Report generated from junctions output from the Trinity CTAT package. For an interactive version of this report see https://igvteam.github.io/igv-reports/examples/example_junctions.html.

### Command line

```
create_reports test/data/junctions/Introns.38.bed
  -genome hg38
  -type junction
  -track-config test/data/junctions/tracks.json
  -info-columns TCGA GTEx variant_name
  -output example_junctions.html
```

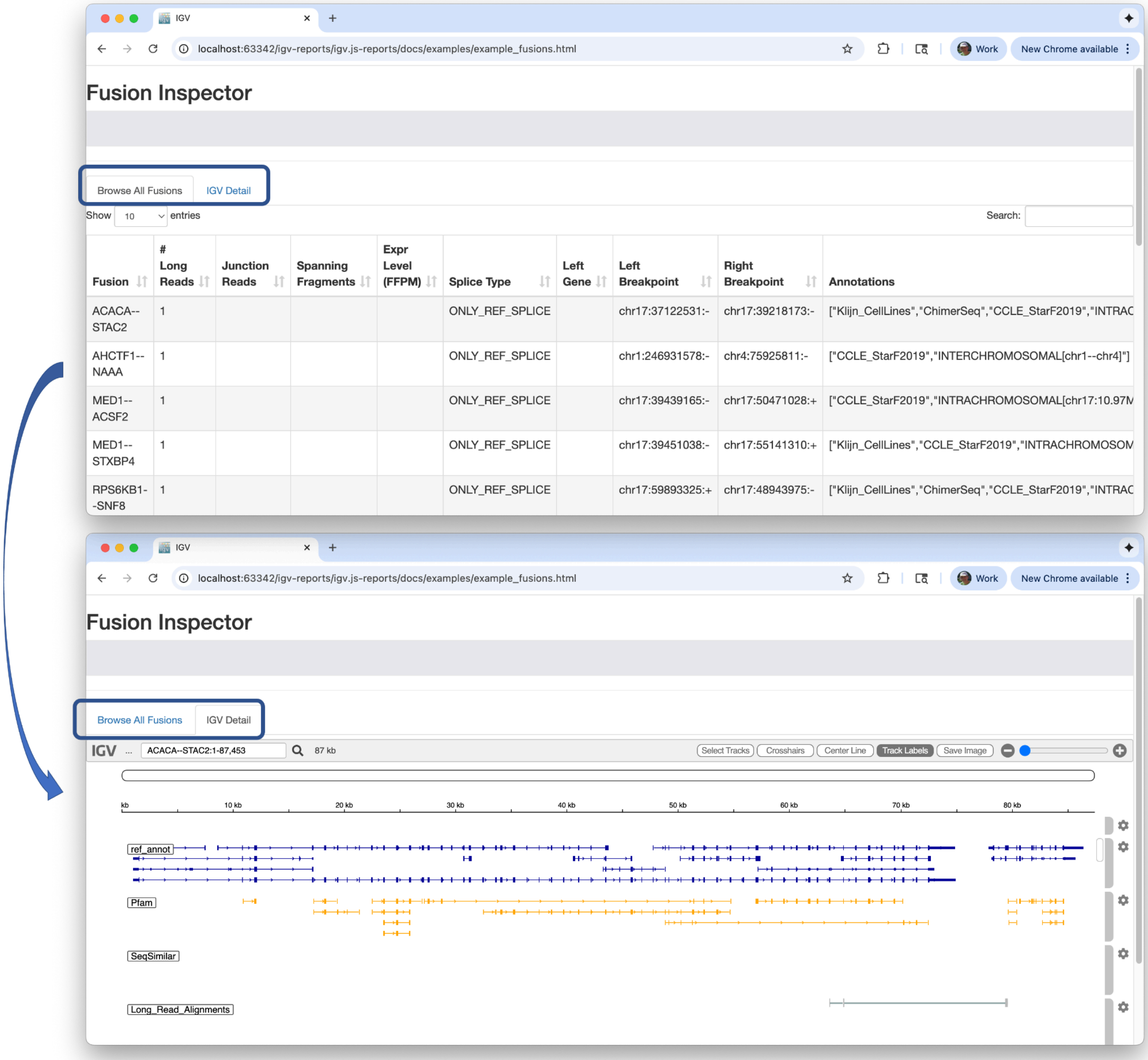


**Supplementary Figure 4.** This report, generated from the Trinity CTAT package's fusion output, features a two-tab interface. The first tab displays a table of fusions, while the second provides an IGV view of the selected fusion row. For an interactive version of this report see https://igvteam.github.io/igv-reports/examples/example_fusions.html.

### Command line

```
create_reports test/data/fusion/igv.fusion_inspector_web.json
  -fasta test/data/fusion/igv.genome.fa
  -template igv_reports/templates/fusion_template.html
  -track-config test/data/fusion/tracks.json
  -output example_fusions.html
```
